# The role of innate immune responses against two strains of PEDV (S INDEL and non-S INDEL) in newborn and weaned piglets inoculated by combined orogastric and intranasal routes

**DOI:** 10.1101/2024.12.20.629601

**Authors:** C. López-Figueroa, E. Cano, N. Navarro, M. Pérez, R. López, K. Skovgaard, H. Vorsholt, P.M.H. Heegaard, J. Vergara-Alert, J. Segalés

**Affiliations:** Unitat mixta d’investigació IRTA-UAB en Sanitat Animal, Centre de Recerca en Sanitat Animal (CReSA), Campus de la Universitat Autònoma de Barcelona (UAB), 08193 Bellaterra, Catalonia, Spain; IRTA, Animal Health, Centre de Recerca en Sanitat Animal (CReSA), Campus de la Universitat Autònoma de Barcelona (UAB), 08193 Bellaterra, Catalonia, Spain; WOAH Collaborating Centre for the Research and Control of Emerging and Re-Emerging Swine Diseases in Europe (IRTA-CReSA), Bellaterra, Barcelona, Spain; Department of Biomedicine and Biotechnology, Technical University of Denmark (DTU), Lyngby, Denmark; Department of Health Technology, Technical University of Denmark (DTU), Lyngby, Denmark; Departament de Sanitat i Anatomia Animals, Facultat de Veterinària, Campus de la Universitat Autònoma de Barcelona (UAB), 08193 Bellaterra, Barcelona, Spain

## Abstract

Porcine epidemic diarrhea (PED) is a severe gastrointestinal disease in swine caused by PED virus (PEDV), leading to significant economic losses worldwide. Newborn piglets are especially vulnerable, with nearly 100% mortality, unlike older pigs. Disease severity also varies depending on the PEDV strain, with non-S INDEL strains being more virulent than S INDEL ones. This study examined early pathogenesis and innate immunity in 1-week-old suckling and 5- week-old weaned piglets (n=8 per age group, 4 per strain) inoculated with S INDEL or non-S INDEL PEDV strains via combined orogastric and intranasal route. Age-matched negative controls (n=3 per age group) were included. Body weight, temperature, and clinical signs were monitored for 48 hours post-inoculation (hpi). PEDV RNA levels were assessed in rectal swabs (RS) at 0 and 48 hpi, while pathological analyses and viral RNA loads were measured in jejunal content and intestinal mucosa. Gene expression of 75 selected antiviral and inflammatory genes were measured in laser capture microdissection (LCM)–derived jejunal samples using microfluidic qPCR at 48 hpi. Suckling piglets showed severe clinical signs, while weaned piglets were mostly asymptomatic at 48 hpi. In general, clinical signs and lesions in suckling piglets were similar, regardless of the PEDV strain. Both viral strains produced comparable viral RNA loads in the small intestine and feces, as well as consistent villous atrophy and fusion across age groups. In LCM-derived jejunal samples, weaned piglets had higher expression of antiviral genes (type I/III interferons, ISGs) and Th1/Th17 pro-inflammatory genes, particularly with the non-S INDEL strain. Conversely, the anti-inflammatory cytokine IL-10 was overexpressed in suckling compared to weaned piglets for both strains. Overall, PEDV-induced intestinal damage, viral replication, and excretion were similar regardless of viral strain or piglet age. The reduced clinical severity in weaned piglets may result from their stronger intestinal antiviral and pro-inflammatory response.

**AUTHOR SUMMARY:** Porcine epidemic diarrhea virus (PEDV) is the causative agent of a major gastrointestinal disease in piglets worldwide, characterized by severe watery diarrhea. The disease is particularly devastating in newborn piglets, especially when caused by non-S INDEL PEDV strains, while weaned piglets demonstrate resistance regardless of the strain. In this study, during the acute infection phase (48 hpi), both highly virulent non-S INDEL and less virulent S INDEL strains caused comparable intestinal atrophy, viral replication in the intestine, and viral loads in feces in both weaned and suckling piglets. However, weaned piglets mounted a robust antiviral response involving type I and III interferons (IFNs) and the induction of Th1- and Th17-related pro-inflammatory responses in the intestinal mucosa. Additionally, interferon-stimulated genes (ISGs) were broadly upregulated across the intestinal mucosa of weaned piglets in response to both PEDV strains. In contrast, suckling piglets exhibited a weaker innate immune response, coinciding with more severe clinical signs. The observed inverse relationship between disease severity and intestinal innate immune activation highlights the potential role of IFNs, ISGs, and pro-inflammatory cytokines in mitigating PEDV severity and underscores their relevance in developing novel pharmacological prevention strategies.

## INTRODUCTION

Porcine epidemic diarrhea (PED) virus (PEDV) is an enteropathogenic *Alphacoronavirus* that causes severe gastrointestinal disease in pigs, characterized by watery diarrhea, severe dehydration, and variable mortality during the acute phase of infection [1]. The disease poses a significant threat to newborn piglets, where mortality rates can approach 100%, while weaned pigs exhibit greater resistance to the virus [2]. The heightened susceptibility in newborn piglets remain poorly understood but it is hypothesized that the immaturity of their gastrointestinal tract and a weaker innate immune response compared to weaned pigs may contribute to the increased mortality [3,4]. Disease severity also varies depending on the PEDV strain, characterized by insertions and deletions (INDEL) in the spike protein S gene, with non-S INDEL strains being more virulent than S INDEL ones. Our previous research in a 5-day-old pig model showed that a highly virulent non-S INDEL PEDV strain caused more severe jejunal damage, leading to increased diarrhea and weight loss compared to S INDEL strains [5], which was consistent with other studies [6–8].

Recent transcriptomic analyses have revealed changes in the expression profiles of numerous innate immunity-related genes following PEDV infection, both in the small intestinal mucosa and *in vitro* cell cultures. Accordingly, the acute onset of diarrhea coincided with differentially expressed genes (DEGs) primarily associated with interferons (IFNs) and proinflammatory cytokines [9–11] in IPEC-J2 cells and intestinal mucosa. However, other essential components of innate immunity, such as those involved in maintaining epithelial barrier integrity, goblet cell function, and the modulation of signaling pathways like MAPK (ERK1/2 and JNK/p38), JAK/STAT, PI3K-AKT/mTOR, and NF-kB, also show significant transcriptional changes upon PEDV infection in both the intestinal mucosa and various *in vitro* cell types. [11–16]. Additionally, biological processes including apoptosis, endoplasmic reticulum (ER) stress, and autophagy are also involved in the innate antiviral response against PEDV [17,18]. These findings emphasize the importance of the mucosal innate immune response in PEDV pathogenesis.

The host’s innate immunity, a key part of mucosal defense, is crucial against porcine coronaviruses (PCoVs) [19]. During PEDV infection, IFNs and proinflammatory cytokines synthesized upon recognition by pattern recognition receptors (PRRs) serve as primary effectors of innate immunity [20–23]. Several studies have demonstrated significant alterations in the expression of proinflammatory cytokines following PEDV infection, including IL-1α, IL-1β, IL-6, IL-8, IL-10, IL-11, IL-12B, IL-18, IL-22, IL-27, CCL2, CCL3, CCL4, CCL5 (RANTES), CXCL2, CXCL10, TNF-α, TGF-β, and IFN-γ [11,24,25]. Notably, certain cytokines such as IL-11, IL- 18, and IL-22 suppress PEDV replication, while IL-6 and IL-8 may contribute to viral persistence [26–30]. On the other hand, both type I (IFN-α/β) and type III (IFN-λ) IFNs are generated upon PEDV infection [20,31,32].

Type III IFNs play a crucial role in protecting epithelial barriers as the IFN-λ receptor (IFNLR1) is predominantly expressed on epithelial cells, whereas IFN-α/β are produced by a number of different cell types [23]. Consequently, type III IFNs are notably more effective in inhibiting PEDV replication at mucosal surfaces [31,33]. Although both type I and III IFNs activate the JAK/STAT pathway to induce expression of interferon-stimulated genes (ISGs) for antiviral defense, their regulatory pathways differ. The NF-κB pathway primarily regulates IFN- λ production, whereas the IRFs pathway predominantly controls type I IFN expression [22].

Additionally, ISGs, which can disrupt viral replication and enhance adaptive immunity, exhibit prolonged activity when induced by type III IFNs, whereas type I IFNs induce a more rapid and transient ISG response [34].

Previous studies investigating the innate immune response to PEDV have primarily utilized cell lines (such as Vero cells and IPEC-J2) or pigs [11,15,24,25,35]. Nevertheless, comparative *in vivo* studies involving piglets of different ages are absent. Therefore, the objective of the present study was to compare early pathogenesis of PEDV infection and analyze the role of the innate immune responses in two different age groups, suckling and weaned piglets, inoculated with either non-S INDEL or S INDEL PEDV isolates.

## MATERIALS AND METHODS

### ETHICS STATEMENT

All animal procedures were approved by the Animal Welfare Committee of the *Institut de Recerca i Tecnologia Agroalimentàries* (CEEA-IRTA, registration number CEEA 86/2022) and by the Ethical Commission of Animal Experimentation of the Government of Catalonia (registration number 11560). The experiments were conducted by certified staff under Biosafety Level 2 (BSL-2) conditions at IRTA-Monells (Girona, Spain).

### VIRAL INOCULA

The European S INDEL PEDV isolate (CALAF 2014, GenBank accession number MT602520.1; PEDV CALAF) and the USA non-S INDEL PEDV isolate (USA/NC49469/2013, GenBank accession number KM975737, generously provided by Dr. J.Q. Zhang and D.M. Madson from Iowa State University; PEDV USA) were used. Both inocula were obtained from jejunal contents and mucosal homogenate of 5-day-old piglets intragastrically inoculated with these PEDV strains [5]. Initially, 200 μL of undiluted intestinal content from each inoculum underwent RT-qPCR to assess the viral load. After vortexing, RNA extraction was carried out using the MagMAX Pathogen RNA/DNA Kit (Thermo Fisher Scientific, Carlsbad, CA, USA, 4462359) following the manufacturer’s instructions. RT-qPCR analysis (Thermo Fisher Scientific, Carlsbad, CA, USA, 4486975) revealed comparable viral RNA loads for PEDV CALAF (Ct value of 12.65) and PEDV USA (Ct value of 11.30) isolates. Viral infectivity titers of each inoculum (10^5^ and 10^6^ tissue culture infectious dose 50% (TCID_50_)/mL, respectively) was extrapolated from the Ct values of the jejunal content using a standard curve, as previously described [36]. The remaining intestinal content along with the intestine homogenate were utilized to prepare the final inoculum for PEDV strains used in this study. For this purpose, original inoculum samples were diluted 1:5 in 1× phosphate-buffered saline (PBS) supplemented with 1% penicillin/streptomycin to a final volume of 90 mL. The mixture was centrifuged for 10 minutes at 800 g and subsequently stored at -80°C until further use.

Additionally, both inocula were confirmed negative for transmissible gastroenteritis virus (TGEV), porcine rotavirus A (PRV-A), porcine reproductive and respiratory syndrome virus (PRRSV), and porcine circovirus 2 (PCV2) using specific RT-qPCR assays (Thermo Fisher Scientific, Carlsbad, CA, USA, with references 4486975, A35751 and QPCV, respectively), as previously described [5].

### EXPERIMENTAL DESIGN

A total of eleven 5-day-old piglets from six sows, which tested negative for PEDV and TGEV in feces examined by RT-qPCR, and with no detectable antibodies against these viruses using commercial ELISAs (Ingenasa, Madrid, Spain, 11.PED.K1 and 11.TGE.K3, respectively), were selected and transported to the experimental farm at IRTA-Monells, Girona, Spain. Sows were also negative for PRRSV and PCV2 in serum confirmed by RT-qPCR and qPCR, respectively. Additionally, eleven 5-week-old piglets, also negative for the same pathogens, were selected and transported to the experimental facilities. Upon arrival, animals were weighed and randomly distributed to three physically separated rooms, with each room designated for a different inoculum: one for each PEDV strain (PEDV CALAF and PEDV USA) and one as a negative control. The experimental design consisted of six groups categorized by age and the type of inoculum administered: (1) 5-day-old piglets inoculated with S INDEL PEDV strain CALAF 2014 (PEDV CALAF 5d), (2) 5-week-old piglets inoculated with S INDEL PEDV strain CALAF 2014 (PEDV CALAF 5w), (3) 5-day-old piglets inoculated with non-S INDEL PEDV strain USA NC4969/2013 (PEDV USA 5d), (4) 5-week-old piglets inoculated with non-S INDEL PEDV strain USA NC4969/2013 (PEDV USA 5w), (5) 5-day-old piglets administrated with PBS (negative control group, CNEG 5d), and (6) 5-week-old piglets inoculated with PBS (negative control group, CNEG 5w). Groups 1–4 contained 4 piglets each, while groups 5 and 6 consisted of 3 piglets each. All had free access to water and were fed every six hours.

On study day (SD) 0, all animals were orogastrically inoculated with 5 mL of the corresponding PEDV isolate or saline solution (CNEG), and intranasally inoculated with 2 mL of the same inocula. Each animal received an equivalent viral titer (10^5^ TCID_50_/mL), irrespective of the PEDV isolate [36].

Pigs were monitored for clinical signs, including rectal temperature and diarrhea, at 0 hpi before challenge, and every 12 h until necropsy at 48 hours post-inoculation (hpi). Diarrhea severity was assessed using a scoring system (0, normal feces; 1, soft and/or pasty feces; 2, watery fecal content, with or without some solid content). Rectal temperatures below 40°C were considered normal, and those ≥40°C were classified as febrile. Body weight was recorded and collected at 0 hpi (before challenge) and at necropsy (48 hpi). RS were collected before challenge (0 hpi) and at necropsy, at 48 hpi. Each swab was placed in 500 μL of minimum essential media, MEM (Thermo Fisher Scientific, Carlsbad, CA, USA, 21090022) supplemented with 1% penicillin/streptomycin (Thermo Fisher Scientific, Carlsbad, CA, USA, 15140122). These samples were used to determine fecal PEDV shedding by RT-qPCR. Viral infectivity, expressed as log10 TCID_50_/mL, was extrapolated from the Ct values obtained from jejunum content using a standard curve, as previously described [36].

### NUCLEIC ACID EXTRACTION AND RT-qPCR/PCR FOR DETECTION OF VIRAL PATHOGENS

Viral RNA/DNA were obtained from feces, RS, and serum samples from sows and piglets (0 and 48 hpi), as well as from jejunal content, intestinal wall (jejunum and colon), nasal turbinate and lungs homogenate from inoculated piglets after 48 hpi. Viral RNA/DNA extraction and the subsequent detection and quantification of various viral pathogens, including PEDV, TGEV, PRV-A, PRRSV and PCV-2 were performed following previously described procedures [5]. Viral load in feces, swabs and tissue macerates were expressed as Ct values.

### DETECTION OF ANTIBODIES AGAINST PEDV AND TGEV

The presence of antibodies against PEDV and TEGV in serum samples from piglets and their respective dams was tested using commercial kits (Ingenasa, Madrid, Spain, 11.PED.K1 and 11.TGE.K3, respectively) as previously described [5]. Results were expressed as mean sample/positive (S/P) ratio. For PEDV, the kit employs an indirect ELISA, with results classified as positive (>0.35) or negative (<0.35). For TGEV, the kit uses a capture blocking ELISA, categorizing results as positive (<1.0008), negative (>1.1676), or doubtful (1.0008–1.1676).

### PATHOLOGICAL ANALYSES AND IMMUNOHISTOCHEMISTRY

All pigs were euthanized at 48 hpi with an intravenous overdose of pentobarbital. Complete necropsies were performed, and tissues including the small intestine (jejunum and ileum), caecum, colon, mesenteric lymph node (LN), nasal turbinates, and lungs were examined for gross lesions by two veterinary pathologists certified by the European College of Veterinary Pathologists, blinded to the treatment groups. Gross findings of the intestinal wall were scored as follows: 0, normal; 1, thin-walled or gas-distended; and 2, presence of both conditions. The contents of the small intestine, caecum and colon were examined and scored: 0, normal; 1, pasty; or 2, watery content.

Samples from the small intestine, colon, mesenteric LN, nasal turbinates, and lungs were fixed by immersion in 10% neutral-buffered formalin and embedded in paraffin. Standard hematoxylin and eosin (H&E) staining was performed. Villus lengths and crypt depths were measured from ten representative villi and crypts in the middle jejunum per animal [17]. The villus-height-to-crypt-depth (VH:CD) ratio was calculated as by dividing the mean villus length by the mean crypt depth.

An immunohistochemistry (IHC) technique to detect the PEDV nucleoprotein (Medgene Labs, Brookings, SD, USA, SD6-29) was performed on selected tissues (jejunum, ileum, colon, mesenteric LN, nasal turbinates, and lungs) following a previously described procedure [5]. Viral antigen presence was semiquantitatively scored from 0 to 3 in intestinal tissues and mesenteric LC using 10 high-power fields (HPF) at 400x magnification, as previously described [5].

### METHACARN-FIXED PARAFFIN-EMBEDDED TISSUE SPECIMENS

To evaluate the mRNA profiles of antiviral and proinflammatory cytokines and detect genomic PEDV, jejunum tissues were fixed by immersion in methacarn at 4°C at 48 hpi to ensure optimal RNA preservation [37]. After 24 hours, tissues were embedded in paraffin, sectioned at 8 to 10 μm thickness and mounted on Arcturus RNase-free polyethylene naphthalate (PEN) membrane glass slides (Thermo Fisher Scientific, Carlsbad, CA, USA, #LCM0522).

Slides were air-dried for 30 minutes at 60°C, deparaffinized using serial-based xylene and ethanol concentrations, and stained with 1% cresyl violet acetate (Sigma-Aldrich, Darmstadt, Germany, #MKCC0238) using standard laboratory procedures. After staining, samples were dehydrated in graded ethanol solutions prepared with RNase-free water, air-dried and stored at -80°C for at least 16 hours. All procedures were conducted under RNase-free conditions using DEPC-treated water (Thermo Fisher Scientific, Carlsbad, CA, USA, #AM9922) and RNaseZap (Thermo Fisher Scientific, Carlsbad, CA, USA, #AM9782) to prevent RNA degradation. After 24 hours, the methacarn-fixed paraffin-embedded (MFPE)-jejunal specimens were processed for laser capture microdissection (LCM) prior to RNA extraction. Immunohistochemistry was also performed on MFPE sections following the procedure described above.

### LASSER CAPTURE MICRODISECTION

For each animal, three consecutive sections from the same MFPE block containing jejunal specimens were cut and processed as described in the previous MFPE specimen section. One section underwent IHC for PEDV nucleoprotein detection to serve as a reference template for the subsequent two sections, which were subjected to LCM.

PEDV-infected enterocytes within the jejunal mucosa identified by IHC in the reference template section, were outlined and micro-dissected using the Leica LMD6500 system (Leica AS LMD; Wetzlar, Germany) with 6.3× magnification, and software settings of 115 power and 10 speed, using the Laser Microdissection 6500 software version 6.7.0.3754. The specific areas microdissected from each cresyl violet-stained MFPE section were then placed into RNase-free 0.5 mL Eppendorf tubes containing 150 µL of buffer PKD from the miRNeasy FFPE Kit (Qiagen, Hilden, Germany, #217504) kept at 4°C. The entire process was conducted under RNase-free conditions to prevent RNA degradation.

### TOTAL RNA ISOLATION AND cDNA SYNTHESIS

Total RNA was extracted from micro-dissected jejunal tissue following the protocol provided by the miRNeasy FFPE Kit (Qiagen, Hilden, Germany, #217504). Briefly, after treatment with proteinase K, RNA was concentrated through ethanol precipitation, purified using RNeasy MinElute spin columns, treated with DNase I for 15 minutes, and finally eluted with 15 µL of RNase-free water, utilizing components from the miRNeasy FFPE Kit. The entire process was conducted under RNase-free conditions. After total RNA extraction, the final RNA concentration was determined using a BioDrop μLite spectrophotometer (Biochrom, Cambridge, UK, #80-3006-55).

RNA samples were stored at -80°C until they were sent to the Department of Health Technology at the Technical University of Denmark (DTU), where cDNA synthesis, preamplification, and qPCR analysis were conducted (see below). cDNA was generated from 250 ng of total RNA/sample, following the manufacturer’s instructions for the QuantiTect Reverse Transcription Kit (Qiagen, Hilden, Germany, #205311). Briefly, RNA (250 ng/sample) was diluted with RNase-free water to a final volume of 12 µL. Subsequently, 2 µL of gDNA buffer was added, and the mixture was incubated at 42°C to remove any genomic DNA contamination. Then, 6 µL of the master mix (containing reverse transcriptase, RT Primer mix, RT buffer, and RNase-free water) was added to each sample, and was then incubated for 15 minutes at 42°C, followed by 3 minutes at 95°C. For further pre-amplification and qPCR analysis, the cDNA was diluted 1:10 in low-EDTA TE buffer.

### TRANSCRIPTOMIC ANALYSES BY MICROFLUIDIC RT-qPCR

qPCR analysis was carried out using the high-throughput BioMark HD system (Fluidigm), and the 96.96 Dynamic Array IFC chips (Fluidigm, South San Francisco, CA, USA, #BMK-M-96.96, which accommodate 96 samples in combination with 96 primer assays). Pre-amplification of cDNA was performed using TaqMan PreAmp Master Mix (Applied Biosystems, Waltham, MA, USA, #4391128), Low-EDTA TE buffer (Avantor, Radnor, PA, USA, #APLIA8569.0500), and Exonuclease I (New England Biolabs, Ipswich, MA, USA, #M0293L), following manufacturer’s instructions. Briefly, 7.6 μL of the pre-amplification master mix was combined with 2.5 μL of the 1:10 diluted cDNA samples, as described above. The samples were then incubated for 10 minutes at 95°C and pre-amplified through 18 cycles of 15 seconds at 95°C and 4 minutes at 60°C. Finally, 4 μL of Exonuclease I master mix was added, followed by incubation for 30 minutes at 37°C, and 15 minutes at 80°C, resulting in a final volume of 14 μL. The pooled undiluted pre-amplified, exonuclease-treated cDNA controls underwent serial dilution in low-EDTA TE buffer (1:2, 1:10, 1:50, 1:250, 1:1250) and were assayed in triplicate to establish relative standard curves and assess primer efficiencies. Additionally, each pre-amplified, exonuclease-treated cDNA sample was diluted 1:10 in low-EDTA TE buffer for later use in qPCR, and each sample was analyzed in duplicate.

The qPCR analysis of all pre-amplified cDNA samples, including both 10x diluted samples and undiluted standard curves, was performed using a mix composed of 2X TaqMan Gene Expression Mastermix (Applied Biosystems, Waltham, MA, USA, #4369016), 20X DNA binding dye (Fluidigm, South San Francisco, CA, USA, #100-0388), EvaGreen Dye 20X (Biotium, Fremont, CA, USA, #31000), and Low EDTA TE-Buffer. The individual assay mixes consisted of Assay Loading Reagent (Fluidigm, South San Francisco, CA, USA) and primer pairs (20 μM for each primer). Following the loading of samples into the wells of the 96.96 Chip (Fluidigm, South San Francisco, CA, USA, #BMK-M-96.96), it was placed in the HX IFC controller for 90 minutes. Subsequently, the chip was transferred to the Biomark HD system for sample quantification. The thermal cycle of the microfluidic qPCR was 600 seconds at 95°C (Hot Start), followed by 35 cycles of 15 seconds at 95°C (denaturation) and 60 seconds at 60°C (amplification and data acquisition). Subsequently, a melting curve analysis was conducted to ensure specific amplification.

### SELECTION OF ANTIVIRAL-RELATED GENES

Seventy-five genes were selected to study the gene expression and transcriptional regulation of the key canonical signaling pathways implicated in antiviral innate immunity and inflammation during PEDV infection. The transcriptomic profiles included analyses of ISGs, type I, II, and III IFNs and their receptors, along with cytokines and chemokines involved in inflammatory responses. Additionally, we examined other innate immune-related genes, such as PRRs and adapter proteins (Aps)/transcription factors (TFs) associated with key PEDV- related signaling pathways. All primer pairs employed in the present study were designed in-house (DTU University, Denmark) and purchased from Sigma-Aldrich (Darmstadt, Germany).

However, the primer sequences specific for PEDV (targeting the Nucleocapsid (N) protein, forward primer at position 27313 and reverse primer at position 27453) were obtained from a previously published study [38]. Detailed information on the qPCR primer sequences and their amplification efficiencies is provided in Supplementary table 1. Whenever possible, primers pairs were designed to span intron/exon borders to minimize the risk of amplifying any contaminating genomic DNA.

Expression data (amplification curves, melting curves, and standard curves) of analyzed genes was collected with the Fluidigm Real-Time PCR analysis software 4.1.3 (Fluidigm Corporation, South San Francisco, CA, USA). The DAG expression software 1.0.5.6. [39] was used to calculate the relative expression of each gene using the relative standard curve method (refer to Applied Biosystems user bulletin #2). Multiple reference gene normalization was performed by using GAPDH, RPL13A and PPIA as endogenous controls. R-squared values, PCR efficiencies and slope value for each assay are described in Supplementary table 2. The up- or down-regulated expression of each gene was expressed as mean fold change (Fc) values after dividing each individual normalized quantity (NQ) value with the calibrator (Supplementary table 3). The mean expression of the three pigs from CNEG 5d and CNEG 5w were used as the corresponding calibrator values to determine the NQ value for each assay.

### STATISTICAL ANALYSES

Statistical analysis of the clinical, pathological, and virological results was conducted using SAS v9.4 software (SAS Institute Inc., Cary, NC, USA). Statistical significance was determined at a level of 0.05 (*p*<0.05). Quantitative variables (weight, temperature, RT-qPCR, viral titer, and VH:CD) were subjected to either parametric tests (ANOVA) or non-parametric tests (Kruskal-Wallis, K-W), depending on the normality and homogeneity of variances assessed through Shapiro-Wilk and Levene tests, respectively. In case of obtaining a significant result, type I error was corrected using Tukey or Bonferroni correction for ANOVA and K-W, respectively. Ordinal categorical variables (diarrhea, lesions, intestinal content, and IHC) were analyzed using a linear model considering binomial distribution, with P-values corrected using Tukey’s Test. However, for comparing different post-inoculation times with hour 0 (in % weight, diarrhea evolution, and temperature), a linear model with binomial distribution was also applied, with P corrected using the Dunnett test in this case. Significance level was set at *p*<0.05.

Logarithmic 10 transformations were applied on Fc values provided by the gene expression data analyses to approach a log normal distribution. Thus, the parametric ordinary one-way ANOVA test corrected with Šídák’s multiple comparisons test was used to compare the means of the logarithmic Fc obtained for each group of animals. Significant gene upregulation or downregulation was considered if they met the criteria of a relative Fc of ≥2- fold or ≤2-fold respectively with *p*<0.05. All the graphs and heatmap were created with Prism version 10 software (GraphPad Software Inc., La Jolla, CA).

## RESULTS

### Newborn piglets experienced more severe disease than weaned piglets, but neither PEDV strain caused fever or mortality after acute infection

By 48 hpi, weight loss was observed in all piglets from the PEDV CALAF 5d and PEDV USA 5d groups compared to their initial weight. This reduction was statistically significant only in the PEDV CALAF 5d group. Specifically, piglets in this group experienced nearly double weight loss compared to those in the PEDV USA 5d group, with reductions of approximately 10% and less than 5%, respectively. In contrast, piglets in both the PEDV CALAF 5w and PEDV USA 5w groups exhibited a statistically significant weight gains ranging from 5 to 10% at 48hpi, similar to the weight gain observed in both CNEG groups (Figure 1A).

**Figure 1.**
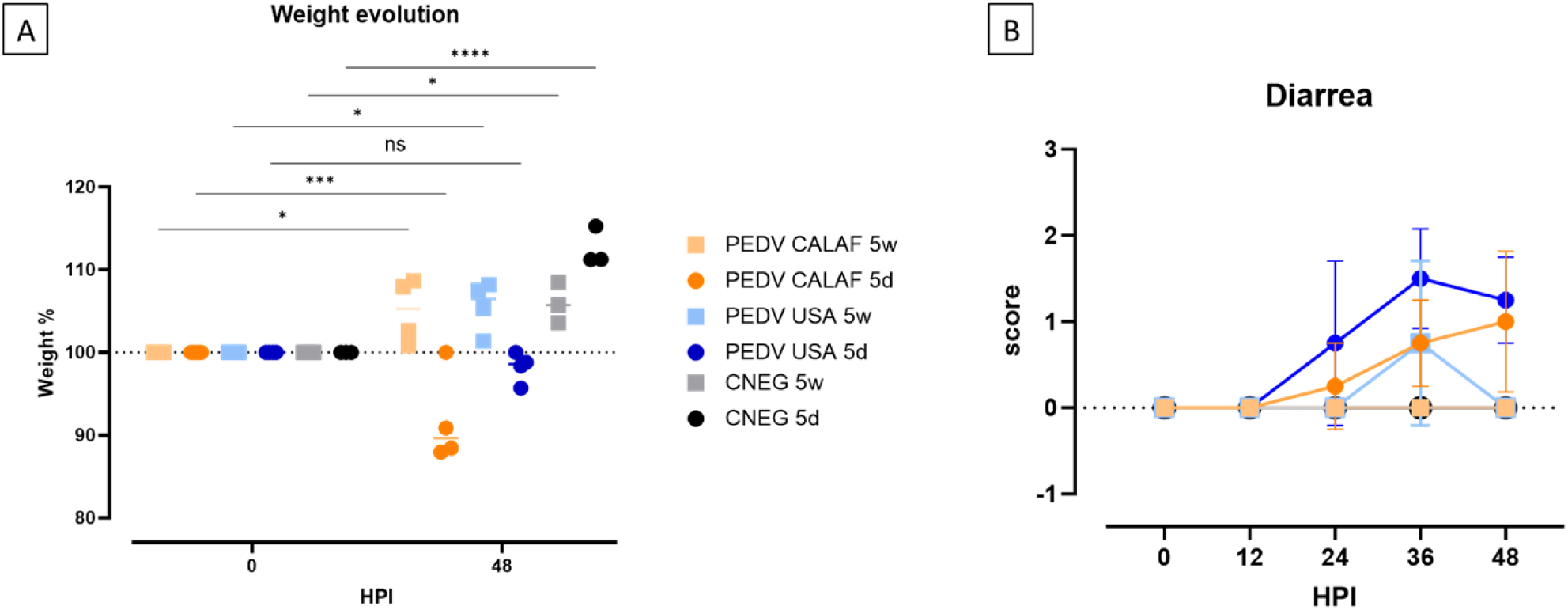
(A) Individual and average weight changes (expressed as a percentage of increase or decrease relative to the initial weight) of the experimental groups from 0 to 48 hpi. (B) Diarrhea evolution over 48 hpi. Diarrhea was scored as no diarrhea (0); pasty (1) and liquid (2) diarrhea. No significant differences between study groups were detected at any HPI. (*, *p*- value < 0.05; ***, *p*-value < 0.001; ****, *p*-value < 0.0001; ns: non-significant).

At 24 hpi, pasty diarrhea was observed in piglets from the PEDV CALAF 5d and PEDV USA 5d groups. However, in the PEDV USA 5d group, diarrhea progressed rapidly to a liquid form by 36 hpi, while in the PEDV CALAF 5d group, liquid diarrhea appeared later, at 48 hpi, in only 2 out of 4 piglets. Among the weaned piglets, pasty feces were observed in the PEDV USA 5w group by 36 hpi in 2 of 4 animals, but no diarrhea was recorded in any animal from the PEDV CALAF 5w group throughout the study period (Figure 1B). Although no statistically significant differences were detected between the groups at any specific time point, a trend toward more severe diarrhea was noted in the PEDV USA 5d group at 36 hpi and in both PEDV USA 5d and PEDV CALAF 5d groups at 48 hpi compared to the CNEG 5d group.

No fever was recorded in any animal throughout the study period.

### Age and strain independence in PEDV-induced small intestinal damage

The summary of individual scores for intestinal wall damage and digestive contents is illustrated in Figure 2.

**Figure 2.**
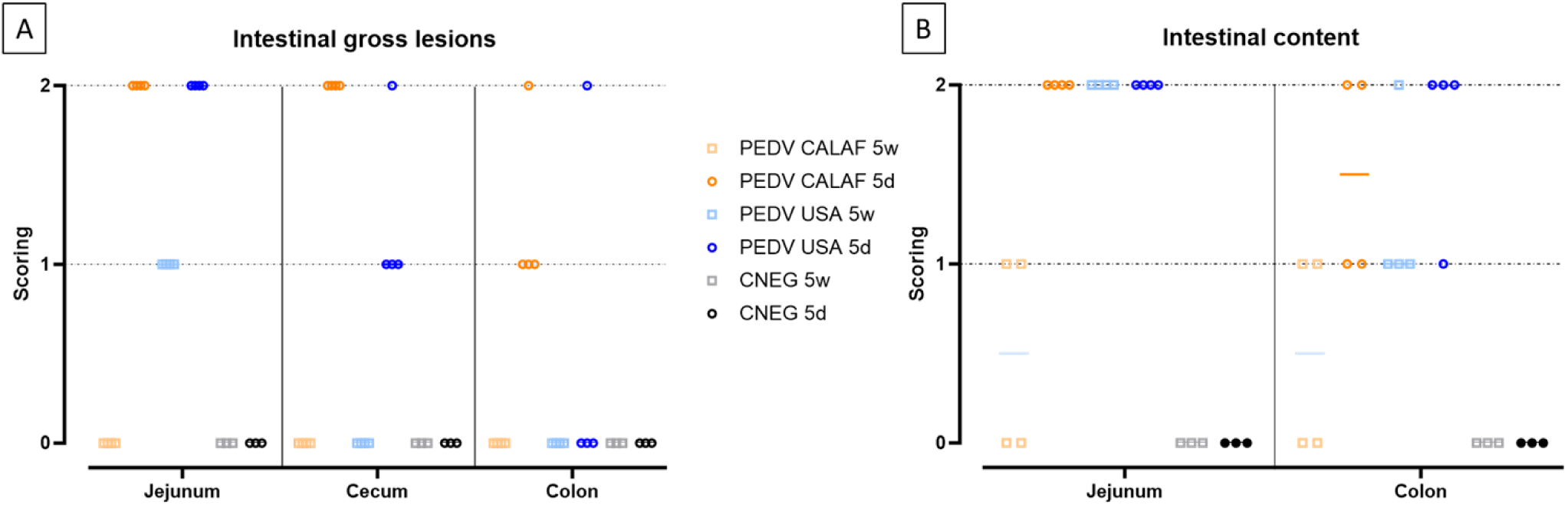
Intestinal lesions (A) and contents (B) in 5-day-old and 5-week-old piglets at 48 hours post-inoculation (hpi) with both strains of PEDV. Intestinal lesion was scored as 0, normal; 1, thin-walled or gas-distended; 2, presence of both. Intestinal content was scored as 0, normal; 1, pasty; or 2, watery content.

At 48 hpi, all newborn piglets in both the PEDV CALAF 5d and PEDV USA 5d groups exhibited thin intestinal walls, gas distension, and liquid contents in the jejunum. In the colon, the PEDV CALAF 5d group showed thin intestinal walls, with 3 out of 4 piglets presenting this condition alone, while 1 piglet exhibited both thin walls and gas distension. Liquid contents in the colon were observed in only 2 piglets from this group. In contrast, only one piglet from the PEDV USA 5d group demonstrated both a thin intestinal wall and gas distension in the colon; nevertheless, 3 out of 4 piglets in this group displayed liquid contents. The remaining animals of both study groups had pasty intestinal contents.

In the PEDV USA 5w group, all piglets exhibited a thin wall and liquid content in the jejunum. Although none of the piglets from this group displayed changes in the colon, three had pasty contents, and one had liquid content. No notable intestinal abnormalities were observed in the PEDV CALAF 5w group; however, two out of four piglets had pasty intestinal contents. Although clear differences observed in intestinal lesions and contents among the groups, these observations did not reach statistical significance.

Histological analysis of jejunum and ileum sections revealed severe, diffuse villus atrophy and fusion in all piglets inoculated with either the PEDV USA or PEDV CALAF strains, irrespective of age (Figure 3A). Enterocytes exhibited vacuolization, cellular hypereosinophilia, and epithelial attenuation, with no evidence of inflammation. These abnormalities were absent in animals administered with PBS. At 48 hpi, intestinal atrophy and fusion were characterized by a notable decrease in villus height (VH), resulting in a reduced VH:CD ratio, which ranged from 1.8 in the PEDV CALAF 5w, PEDV USA 5d, and PEDV USA 5w groups to 2.5 in the PEDV CALAF 5d (Figure 3B). This decrease in VH:CD was statistically significant when compared to the negative controls, which had VH:CD ratios of 4 and 6 in the CNEG 5w and CNEG 5d groups, respectively. No lesions were identified in the colon, nasal turbinates, or lungs of any piglet inoculated with either PEDV isolate or in the negative control piglets.

**Figure 3.**
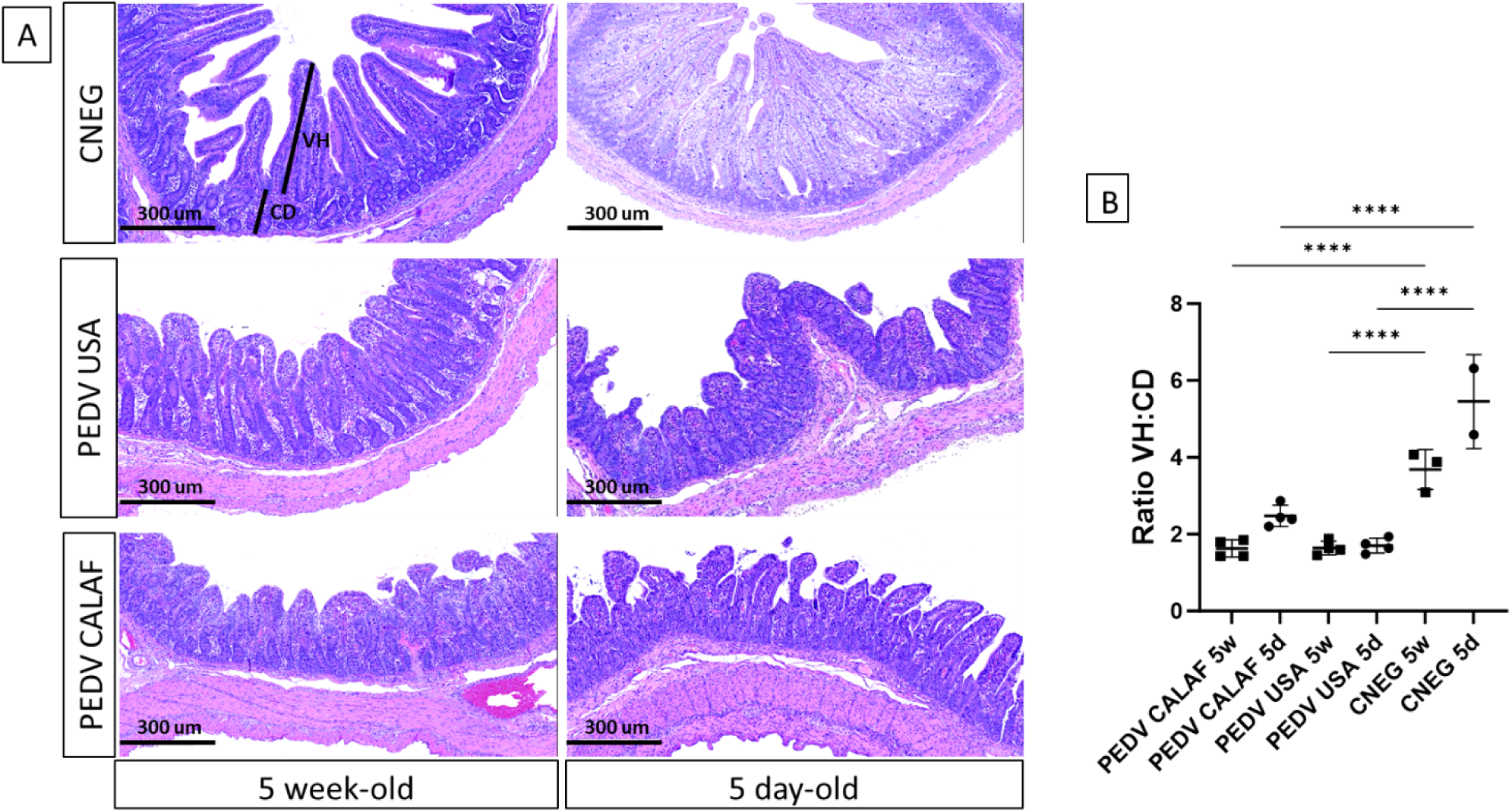
(A) Jejunum at 48 hpi from 5-day-old and 5-week-old piglets from the negative control (CNEG), PEDV USA and PEDV CALAF groups stained with Hematoxylin and Eosin (H&E). Similar severe villous atrophy and fusion was observed in both PEDV groups regardless of age compared to their respective CNEG. (B) The extent of intestinal damage was assessed using the villus height/crypt depth ratio (VH:CD) (**, p-value < 0.01; ****, p-value < 0.0001).

PEDV antigen, detected by IHC, exhibited diffuse and granular staining patterns, primarily located within the enterocytes of the jejunum and ileum in all PEDV-inoculated piglets, regardless of PEDV strain and age (Figure 4A). Viral antigen was widely distributed in the apical portion of the intestinal villi, corresponding to areas of villus atrophy and fusion, and occasionally observed in the crypts of Lieberkühn and in interstitial mononuclear cells dispersed throughout the lamina propria (Figure 4A, indicated by arrows and arrowhead).

**Figure 4.**
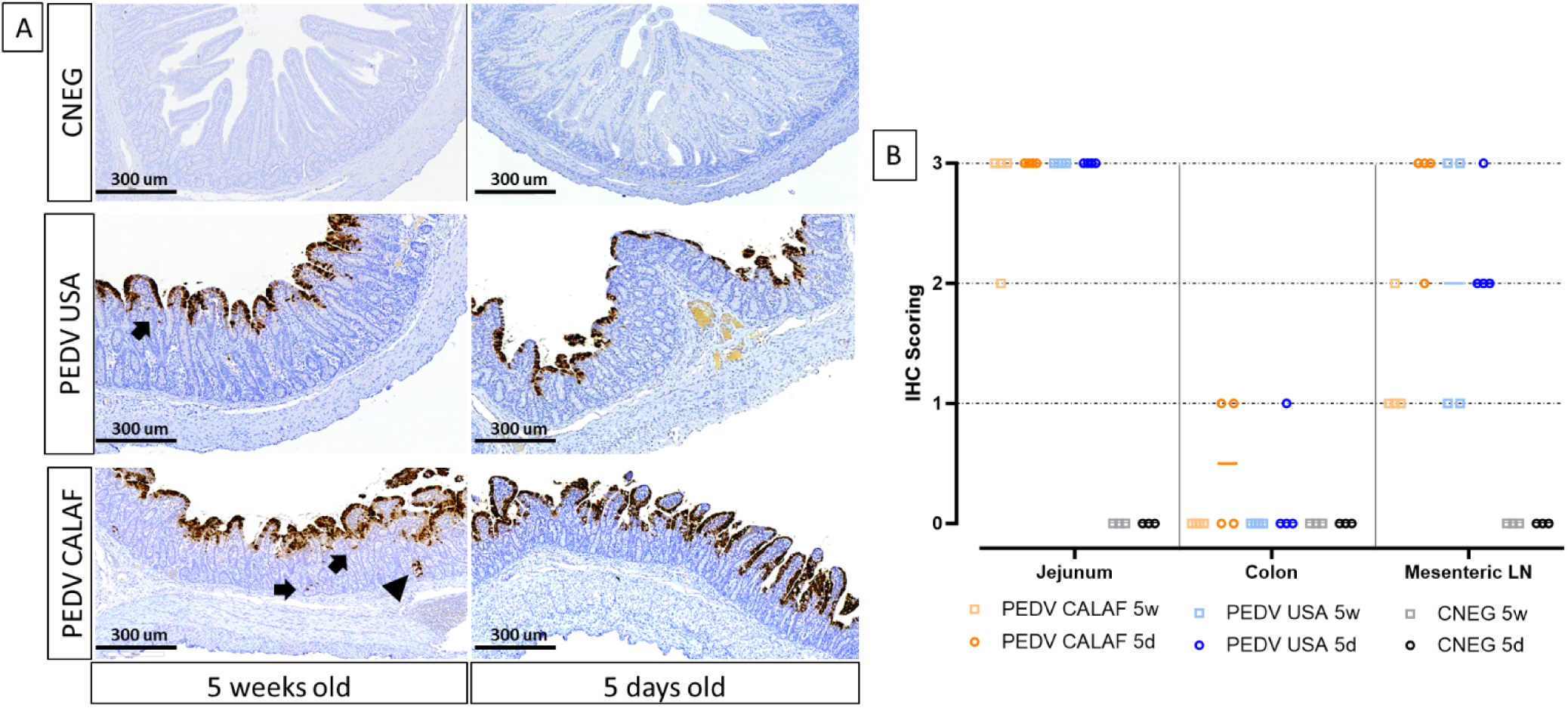
(A) Detection of PEDV by IHC in sections of jejunum of 5-day-old and 5-week-old piglets at 48 hpi. PEDV was similarly detected in the jejunum of the different study groups, regardless of age. PEDV antigen was distributed widely in the cytoplasm of enterocytes from the apical part of atrophic and fused villi. IHC for PEDV detection, hematoxylin counterstain. Detection of PEDV in occasional intestinal crypts (arrowhead) and mononuclear cells of the lamina propria (arrow). (B) Individual IHC scoring of PEDV in jejunum, colon, and mesenteric LN at 48 hpi.

While the jejunum contained the highest amount of PEDV antigen in all animals from inoculated groups, PEDV antigen in the colon was sparsely detected in a few enterocytes in only 2 out of 4 piglets from the PEDV CALAF 5d group and in 1 out of 4 piglets from the PEDV USA 5d group (Figure 4B). Additionally, PEDV antigen was detected in dendritic-like cells within the mesenteric LN. The detection levels in these cells were comparable between the PEDV CALAF 5d and PEDV USA 5d groups, with both showing a higher amount antigen presence compared to that of 5-week-old piglets. Among the latter, a higher amount of PEDV antigen was observed in the PEDV USA 5w group compared to the PEDV CALAF 5w one. Statistical analysis, however, did not reveal significant differences among the study groups.

No PEDV antigen was identified in sections of the nasal turbinate or lungs in any PEDV- inoculated piglet inoculated, or in intestinal segments of PBS-administered piglets.

### non-S INDEL and S INDEL PEDV isolates had similar loads in small intestine and feces in newborn and weaned piglets

The amount of PEDV RNA was analyzed by RT-qPCR in RS at 0 and 48 hpi, as well as in samples from jejunal and colon mucosa, jejunal content, nasal turbinate, and lung macerates at 48 hpi from all piglets.

At 0 hpi, RS from all groups tested negative for PEDV. However, by 48 hpi, viral RNA was detected in fecal samples (Figure 5A), intestinal wall sections (Figures 5B and 5C), and jejunal content (Figure 5D) in all PEDV-inoculated piglets, with comparable levels of viral RNA across groups. No viral RNA was detected in the CNEG groups. The differences in viral RNA levels observed between the PEDV-inoculated groups were not statistically significant.

**Figure 5.**
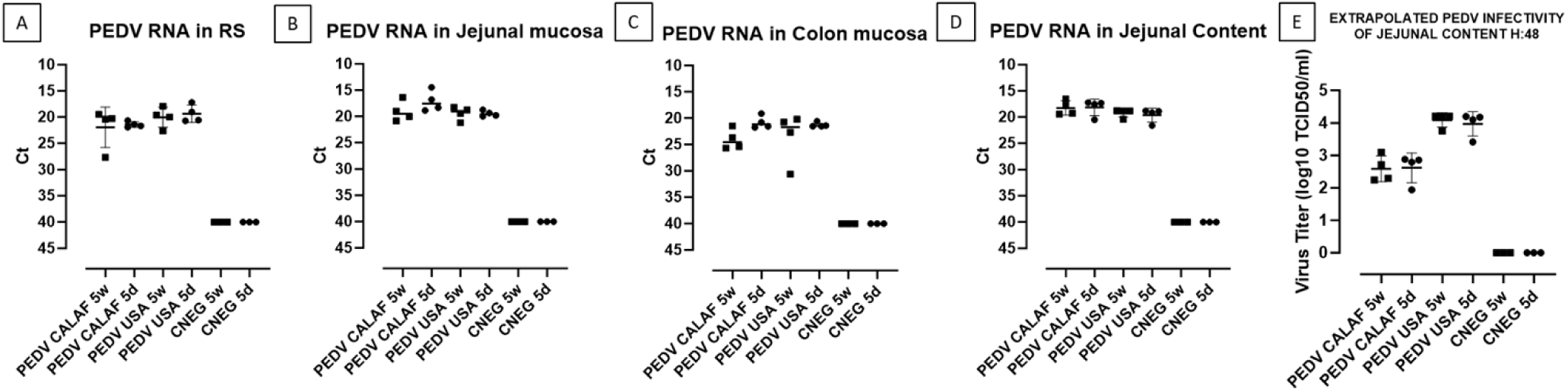
PEDV loads detected in rectal swabs (RS) (A), jejunal (B), and colonic (C) mucosa, and jejunal content (D) at 48 hpi. Fecal viral infectious titer (VT), expressed as inferred log10 TCID_50_/ml, was calculated from Ct values of jejunal content at 48hpi (E).

Additionally, all inoculated piglets shedded infectious virus, as evidenced by viral titration in the jejunal content (Figure 5E). The levels of infectious PEDV were similar between 5-day-old and 5-week-old piglets for each PEDV isolate. However, a higher viral titer was observed in both PEDV USA age groups (10^4^ TCID_50_/mL) compared to both PEDV CALAF age groups (10^2.5^ TCID_50_/mL), although the difference was not statistically significant.

PEDV RNA was detected in the nasal turbinates of all animals in the PEDV CALAF 5w group, but only in a few animals from the other groups, with Ct values ranging from 33 to 38 (Figure 6A). Also, in the lungs (Figure 6B), PEDV RNA was found in all piglets from the PEDV CALAF 5w group and in only one piglet from the PEDV USA 5w group. Interestingly, no PEDV RNA was detected in the lungs of 5-day-old piglets inoculated with either PEDV isolate or in those from the control groups. Nonetheless, no statistically significant differences were observed between the study groups or their respective negative controls.

**Figure 6.**
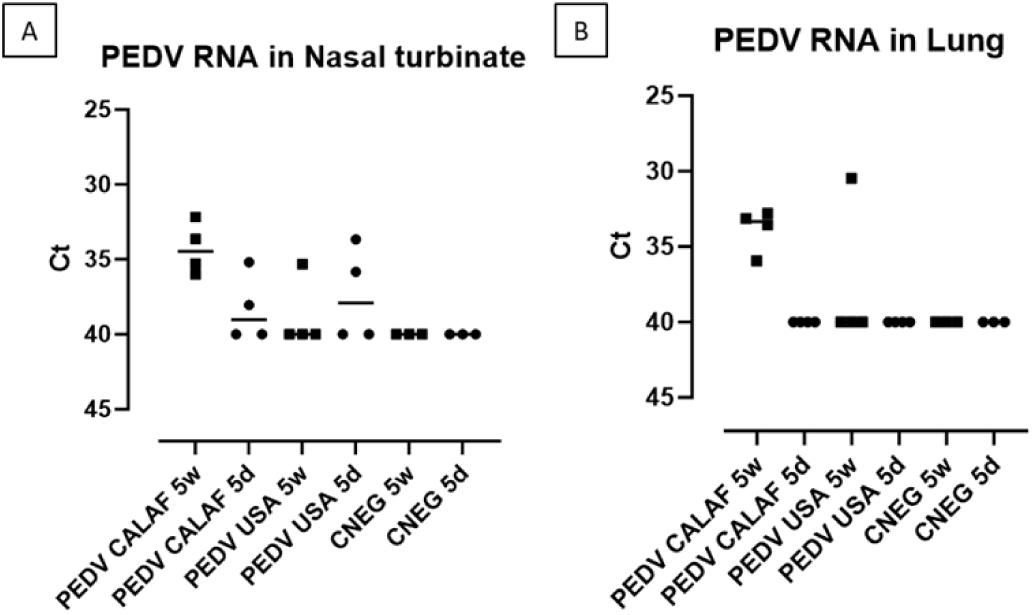
PEDV loads (measured as Ct values) detected in nasal turbinate (A) and lung (B) at 48 hpi of suckling (5d) and weaned (5w) piglets inoculated with PEDV CALAF and PEDV USA strains.

All viral inocula and all samples tested negative for TGEV and PRV-A, as determined by RT-qPCR. The detection limit for all RT-qPCR was set at a Ct value of 40.

### Both PEDV strains actively replicated in the jejunum of newborn and weaned piglets

PEDV loads in enterocytes from micro-dissected MFPE intestinal samples were quantified using specific primers for the N protein employing a microfluidic quantitative PCR assay. The limit of detection for the PEDV N gene was set at Ct = 25.26, ensuring accurate quantification with an average Tm value of 79.45, which confirmed the specificity of the amplification. Notably, the amount of PEDV N RNA levels in the jejunal sections were twice as abundant in the CALAF 5d group (NQ: 2.55) compared to the other groups, which had an average NQ of 0.5. This observation contrasted with the PEDV viral load from the jejunal mucosa macerates and the IHC results, which did not show such a pronounced difference (Supplementary table 4).

### Weaned piglets mount a stronger antiviral (IFN I/III and ISGs) jejunal response against PEDV USA and PEDV CALAF isolates compared to newborn piglets

Normalized quantity (NQ) values for each assay were converted into Z-scores and presented in a heatmap (Figure 7) to illustrate gene regulation patterns in the intestinal mucosa of suckling and weaned piglets. Overall, at 5 days of age the PEDV USA group showed upregulation at 48 hpi in jejunal tissue of more IFN and ISG related genes than the PEDV CALAF group (Figure 7). When infected at weaning, both groups showed significant upregulation of both antiviral and proinflammatory gene expression. Overexpressed genes in suckling and weaned pigs at 48 hpi exhibited a relative expression increase of more than two-Fc compared to the CNEG animals (Figure 8). Although numerous differences in gene transcription profiles between age groups achieved statistical significance (*p*<0.05), others did not, possibly because of the limited sample size and substantial individual variability in gene expression levels within the groups.

**Figure 7.**
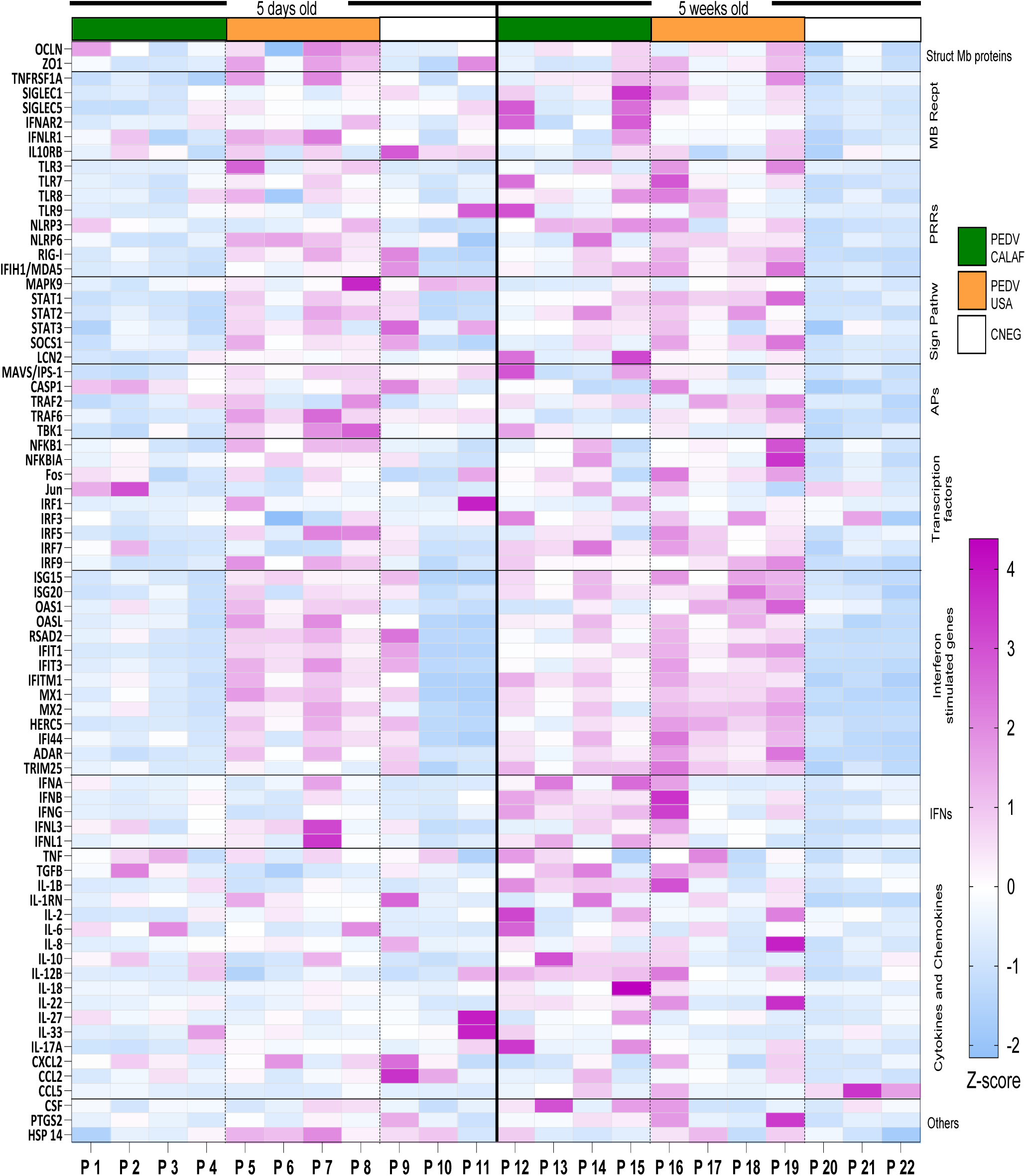
Transcriptional expression of innate immune response-related genes in the jejunum at 48 hours after PEDV CALAF and PEDV USA infection in 5-day-old and 5-week-old piglets, respectively. Each column represents an individual piglet (P1 to P22) within the different age (right and left) and inoculation group (PEDV CALAF, green color; PEDV USA, orange color; CNEG, white color). The resulting heatmap shows color variations corresponding to Z-score; blue indicates decreased gene expression, while pink represents increased expression, relative to piglets in the CNEG groups. Mb Recpt: Membrane receptors; PRRs: Pattern recognition receptors; Sign Pathw: Signaling pathways; APs, adaptor proteins; IFNs: Interferons.

**Figure 8.**
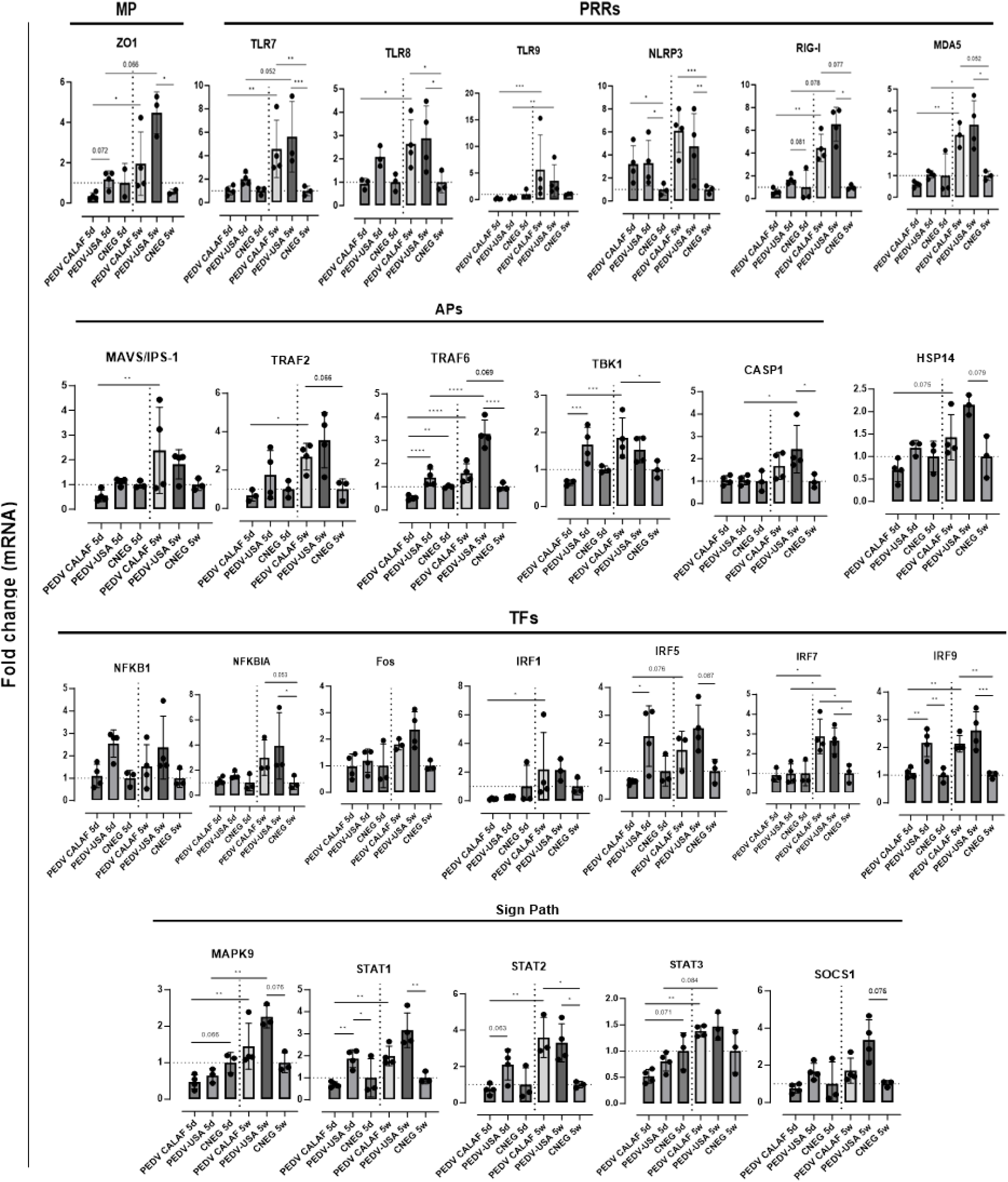
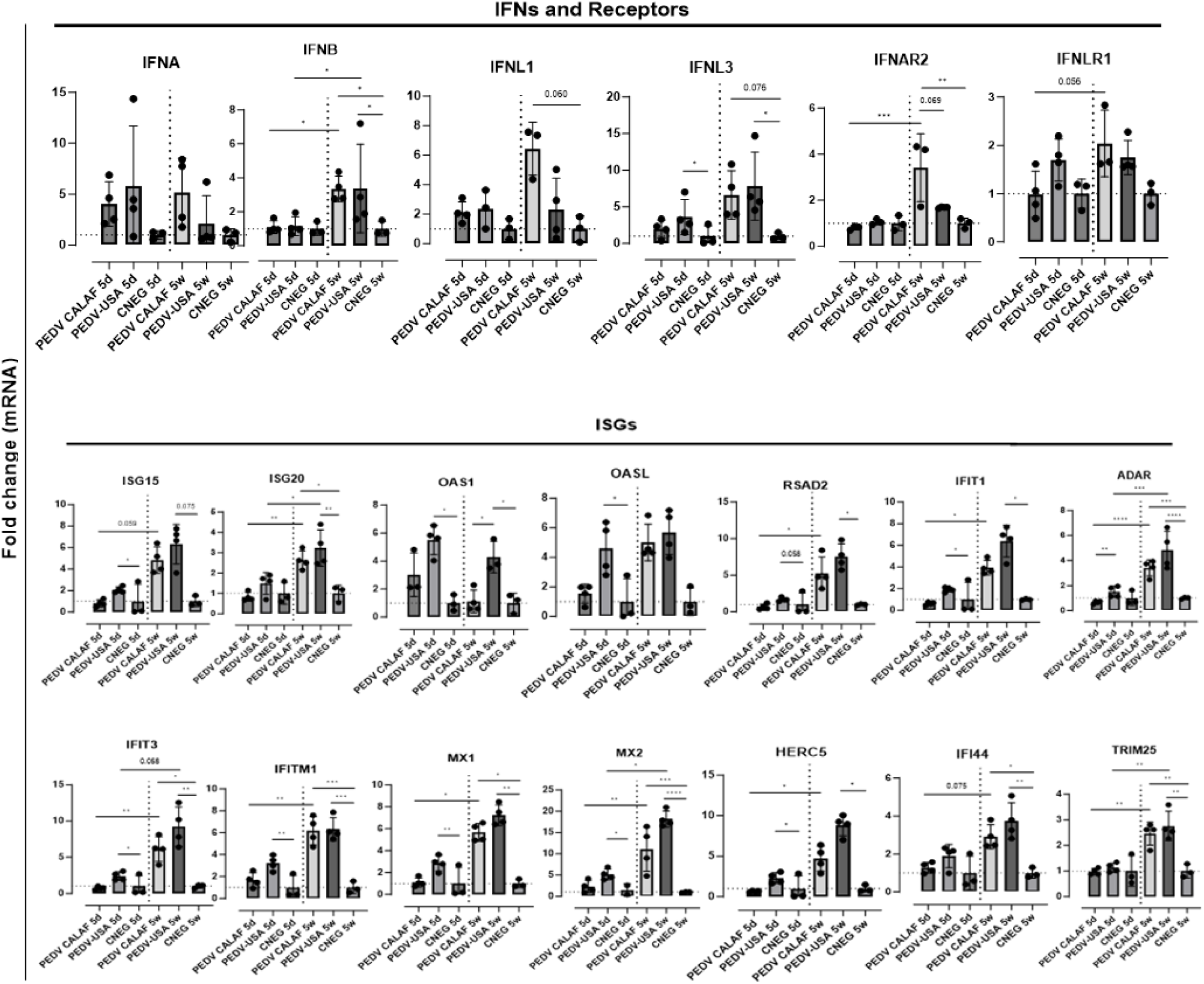
Average fold change of upregulated genes involved in antiviral innate immune response in the jejunum of piglets (5-day-old and 5-week-old) infected with PEDV CALAF and PEDV USA strains, compared to piglets in the CNEG groups. Genes were classified in different categories regarding their main function: MP: Structural membrane proteins; PRRs: Pattern recognition receptors; Sign Path: Signaling pathways; APs: adaptor proteins; TFs: Transcription factors; IFNs: Interferons; ISGs: Interferon stimulated genes. (*, p-value < 0.05; **, p-value < 0.01; ***, p-value < 0.001; ****, p-value < 0.0001). Dashed lines represent baseline expression levels from negative control animals, used as calibrators to calculate the NQ values for each assay.

A consistent pattern of jejunal gene expression was observed across all levels of innate immune-related downstream pathways, from PRRs to IFNs and ISGs, in response to infection with PEDV (Figure 8). Genes coding for IFN-β, IFN-λ1, and particularly IFN-λ3 were significantly upregulated in all piglets from PEDV CALAF 5w and PEDV USA 5w groups relative to piglets in the CNEG groups. In contrast, in 5-day-old piglets, IFN-λ1 and IFN-λ3 were slightly although not significantly upregulated in response to the PEDV USA isolate, and to a lesser extent to the PEDV CALAF strain. Notably, IFN-β remained unaltered in newborn piglets compared to both groups of weaned piglets. In contrast, IFN-α showed similar upregulation across all study groups compared to the CNEG groups, with no significant differences between them (supplementary figure 1).

A similar transcription profile was noted for most PRRs, such as cytoplasmic RNA sensors (RIG-I, MDA5, and NLR, particularly NLRP3), viral endosome double- and single-stranded RNA sensors (TLR3 and TLR7/8, respectively), and cytoplasmic DNA sensors (TLR9). This pattern extended to most PRR-related APs (such as TRIM25, MAVS/IPS-1, TRAF2/6 and TBK1) and TFs (such as IRF1, IRF7 and NF-κB). However, unlike the other TFs, IRF3, which is crucial for the induction of type I IFNs (IFN-α and IFN-β), showed no change in transcription in any of the infected groups. In addition, the relative expression of genes encoding IFN receptors (IFNAR2 and IFNLR1), TFs involved in ISG synthesis (IRF9, STAT1 and STAT2), antiviral ISGs (ISG15, ISG20, OASL, RSAD2, MX1, MX2, IFIT1, IFIT3, IFITM1, HERC5, IFI44 and ADAR) and IFNs/ISGs and proinflammatory regulators (STAT3 and SOCS1) increased similarly in the PEDV CALAF 5w and PEDV USA 5w groups. In 5 day-old-piglets the expression of none of these genes was induced by CALAF PEDV infection while they were moderately induced by PEDV USA.

Furthermore, HSP14 was significantly upregulated in piglets from PEDV USA 5w group and moderately upregulated in those from the PEDV CALAF 5w group, compared to 5-day-old piglets. Nevertheless, the relative expression of HSP14 in the PEDV USA 5d group was similar to that of the control group, while it was downregulated in the PEDV CALAF 5d one.

Overall, weaned piglets exhibited a pronounced intestinal antiviral response against both non-S INDEL (PEDV USA) and S INDEL (PEDV CALAF) strains of PEDV, contrasting with the relatively weak response in newborn piglets. Interestingly, newborn piglets exposed to the non-S INDEL PEDV strain showed higher levels of gene transcription compared to the S INDEL one.

In all control non-infected animals, except for one piglet (P9), all gene transcripts were at baseline levels and the mean expression of the three pigs were used as calibrator values to determine the NQ value for each assay (Supplementary table 3). In that particular piglet, the relative expression of most antiviral and proinflammatory genes was similar to that of PEDV USA 5d group, although no evidence of PEDV infection based on RT-qPCR or IHC. This suggest that observed response was unrelated to PEDV infection.

### Both non-S INDEL and S INDEL PEDV strains modulated the MAPK signaling pathway in weaned piglets but not in newborn piglets

At 48 hpi, five-week-old piglets from the PEDV CALAF and PEDV USA groups exhibited a significant increase in the expression of genes involved in cell survival such as the MAPK9 and Fos (a subunits of the MAPK-dependent activator protein 1 (AP-1) complex) gene compared to the response of infected 5-day-old piglets (Figure 8). This transcriptional pattern correlated with the expression of TRAF6 and TRAF2, APs critical for activating both the NF-κB and MAPK/JNK-p38 singaling pathways. Although the increase following infection in MAPK9 gene expression was statistically significant when comparing 5-day-old and 5-week-old piglets, the difference in Fos gene expression did not reach statistical significance.

### Junctional membrane and pyroptosis-related genes were upregulated in weaned piglets but not in newborn piglets inoculated with both PEDV strains

To evaluate putative cellular changes following PEDV infection, we examined the gene expression of junctional membrane proteins that are critical for enterocyte membrane integrity, such as zonula occludens-1 (ZO-1) and occludin (OCLN). In weaned piglets from both PEDV groups, there was a slight increase in OCLN transcriptional expression, while the ZO-1 gene was significantly upregulated post-infection (Figure 8). These responses were not observed in newborn piglets. Additionally, the expression pattern of CASP-1 gene, associated with pyroptosis, mirrored the significant upregulation observed in ZO-1 in weaned piglets.

### Weaned piglets mounted a proinflammatory intestinal response mediated by Th1 and Th17 cytokines, and prostaglandins against both PEDV strains, whereas this response was suppressed in newborn piglets

In response to PEDV infection, the pro-inflammatory transcription factor NF-κB was slightly upregulated across all study groups, particularly in piglets inoculated with the PEDV USA strain, regardless of age (Figure 8), though this difference was not statistically significant. Conversely, the NF-κB inhibitor (NFκBIA or IκBα) was notably upregulated in PEDV CALAF and PEDV USA-inoculated weaned pigs compared to newborn piglets, where no changes were observed in comparison to CNEG 5d group.

Most genes coding for pro-inflammatory cytokines (IFN-γ, TNF-α, IL-27, GM-CSF, and IL-17A) produced by Th1 and Th17 responses were similarly upregulated in weaned piglets inoculated with either PEDV CALAF or PEDV USA (Figure 9). Similarly, the intestinal expression of the immune regulators IL-2 and TGF-β were increased in weaned piglets from both PEDV groups. In contrast, these cytokines remained at baseline levels in newborn piglets, similar to the CNEG 5d group. Other pro-inflammatory factors like IRF5, IL-12B, IL-1RN, IL-22, IL-8, and IL- 33 significantly increased at 48 hpi in weaned piglets compared to newborn piglets after infection. Notably, the expression of IL-1RN was remarkably high and restricted to all 5-week-old piglets. Moreover, CASP-1, IL-1β, and IL-18, key components of the NLRP3 inflammasome and NK cells and CD8+ cytotoxic T lymphocytes (CTLs) promoters (IL-18), exhibited significant mRNA upregulation in weaned piglets compared to newborn piglets. Furthermore, the prostaglandin gene (PTGS2) was upregulated in PEDV CALAF 5w and PEDV USA 5w groups, with no transcriptional activity observed in newborn piglets.

**Figure 9.**
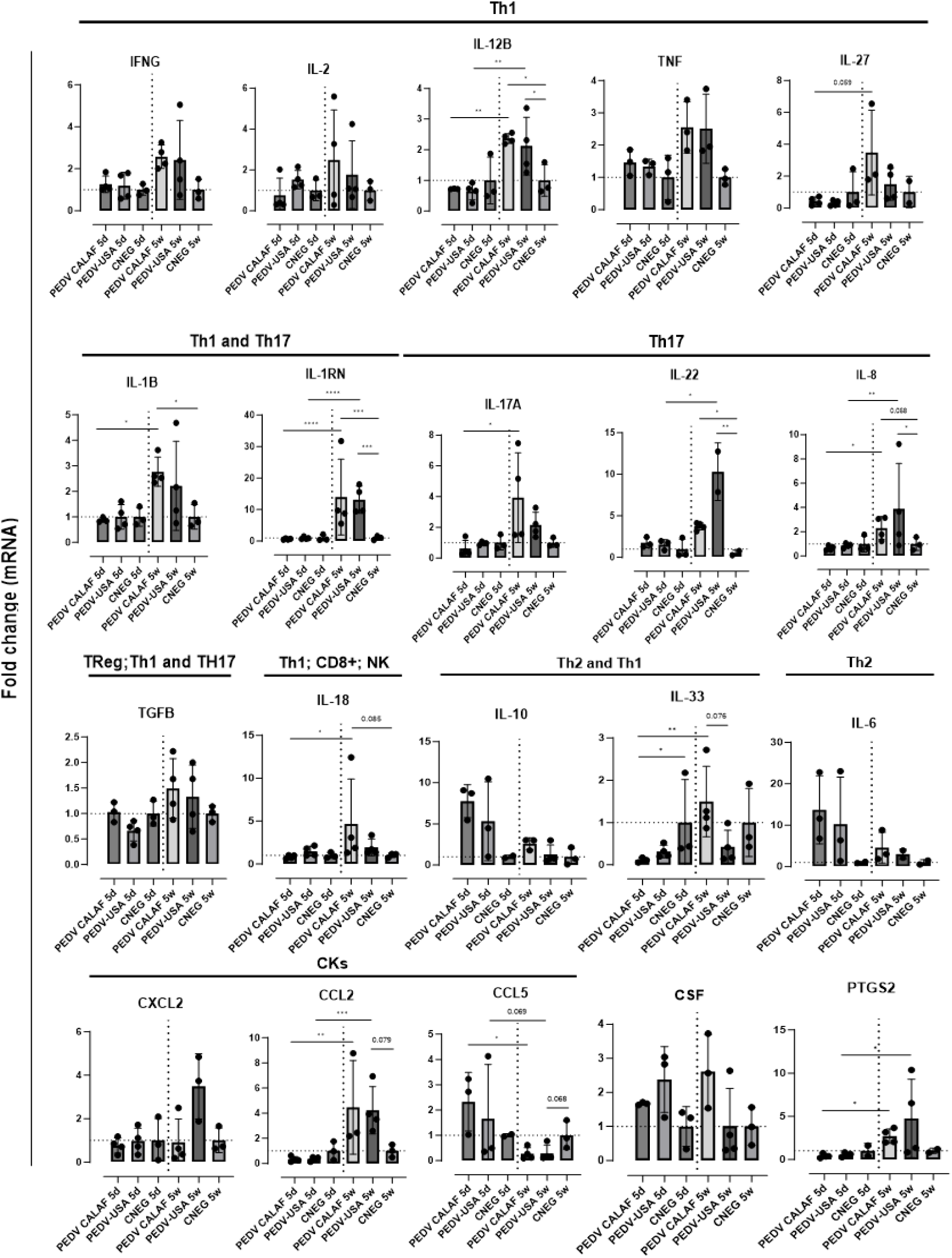
Average fold change of genes upregulated after infection involved in the inflammatory response in the jejunum of piglets (5-day-old and 5-week-old) infected with PEDV CALAF and PEDV USA strains. Genes were classified in Th1, Th2 and Th17 phenotypes as well as involved in Treg, natural killer (NK) cells and CD8+ (cytotoxic T-Lymphocytes) cell activation. CKs: Chemokines; CSF: Colony Stimulating Factor; PTGS: prostaglandin. (*, p-value < 0.05; **, p-value < 0.01; *** > p-value < 0.001; ****, p-value < 0.0001). Dashed lines represent baseline expression levels from negative control animals, used as calibrators to calculate the NQ values for each assay.

The chemokine CCL2 was similarly upregulated in 5-week-old piglets from both PEDV CALAF and PEDV USA groups, while CXCL2 was only overexpressed in the PEDV USA 5w group (Figure 9). None of these chemokines showed altered gene expression in 5-day-old piglets from any group. Conversely, CCL5 gene expression was induced in newborn piglets from both PEDV CALAF and PEDV USA groups, compared to weaned piglets, in which its expression was downregulated relative to the control.

Similar to CCL5, Th2-related cytokines such as IL-6 and the pro-inflammatory antagonist IL-10 exhibited higher expression levels in 5-day-old piglets inoculated with both PEDV CALAF and PEDV USA, whereas their expression was marginal in all weaned piglets (Figure 9).

## DISCUSSION

The impact of PEDV infection depends on host factors like age, innate immune response, and cellular pathways related to cell damage and survival [1,18,23]. Virus-specific factors such as PEDV strain type (non-S INDEL or S INDEL), inoculum dose, and herd immunity status also play key roles in disease outcome [1,6,7,40]. The present study compared the transcriptomic responses between newborn (5-days-old) and weaned (5-week-old) piglets after inoculation with either the European S INDEL (CALAF) or North-American non-S INDEL (USA) PEDV isolates to explore age-dependent immune and inflammatory responses.

In line with previous literature [1,2], the clinical outcomes in the present study were age and PEDV strain-dependent. Newborn piglets suffered from severe diarrhea, weight loss, and pronounced intestinal lesions, whereas weaned piglets developed less severe symptoms, with moderate to severe gross intestinal lesions and pasty diarrhea. Although the non-S INDEL strain caused earlier onset and more severe diarrhea in newborn piglets compared to the S INDEL strain, as previously reported [6–8], the S INDEL group experienced more significant body weight loss. However, contrary to expectations, no mortality was observed, even among newborn piglets infected with the more virulent non-S INDEL strain—a surprising deviation from established literature [2]. Furthermore, pathological and virological outcomes, including intestinal damage (villi atrophy and fusion), PEDV distribution in the jejunum, replication, and virus shedding in feces were similar across all groups. These findings contradict earlier literature, which suggested that these parameters are more pronounced in newborn piglets, especially when infected with the non-S INDEL strain [6–8].

These observations suggest different explanations for the lack of evident clinical signs in weaned piglets despite viral replication, excretion, and intestinal damage. First, clinical signs in weaned piglets may appear later than in newborns [1], suggesting that diarrhea might have developed if the study would have been extended. Additionally, resistance to PEDV infection may be affected by differences in intestinal development between newborn and weaned piglets, as noted in previous studies on the small intestine [3,7]. In addition to the small intestine, this study also found gross lesions and PEDV antigen in the large intestine, but only in newborn piglets. Therefore, the more developed, lesion-free colon in weaned piglets may help mitigating small intestinal diarrhea, preventing overt clinical signs. However, further studies are needed to explore other factors influencing PEDV pathogenesis, such as differences in intestinal microbiota and anatomical or physiological factors, including key functional membrane receptors in small and large enterocytes across different piglet ages.

Moreover, the upregulation of junctional membrane protein genes (ZO-1) in jejunal enterocytes from weaned piglets after PEDV inoculation indicates an adequate response to cellular disruption, aligning with previous studies [13,41]. This suggests that weaned piglets have better intestinal regeneration capabilities, which mitigate PEDV-induced enterocyte damage and explain the lack of evident clinical signs. Conversely, newborn piglets show unchanged ZO-1 gene expression, suggesting either limited intestinal regeneration or PEDV- mediated transcriptional inhibition, as previously reported [7,18].

The efficiency of the innate immune response is a key factor in determining clinical outcomes of PEDV infection in newborn versus older piglets [3]. In this context, the integrity of the mucosal barrier, a vital component of innate immunity, is crucial for defense against pathogens such as PCoVs [16,18,42]. Therefore, special attention was given here to characterize innate immune responses in the jejunum during the acute phase of PEDV infection (48 hpi). Microfluidic qPCR analysis on LCM-derived jejunal enterocytes with high viral loads showed distinct gene transcription patterns for antiviral and inflammatory responses between neonatal and weaned piglets, correlating with the observed age-dependent clinical outcomes. These findings underscore that in this study the development of effective innate immune and inflammatory responses is influenced by age rather than by the specific strain of PEDV.

Regarding the antiviral response, weaned piglets similarly upregulated the intestinal gene expression of IFNs and ISGs against both PEDV strains at 48 hpi. Conversely, these genes were weakly expressed in newborn piglets, a fact apparently associated with severe clinical outcomes. However, antiviral IFNs and ISGs were more upregulated in newborn piglets inoculated with the more virulent non-S INDEL strain compared to the S INDEL strain. This suggests that the S INDEL strain may modulate the expression of these genes to evade the host immune response, promoting viral replication, which aligns with the higher viral load (PEDV N mRNA) observed in the PEDV CALAF 5d group [22,43]. Additionally, the resultant gene transcription pattern in newborn piglets infected with non-S INDEL PEDV strains may explain the lack of mortality, highlighting the antiviral role of IFNs and ISGs in controlling PEDV replication.

Based on the obtained results, Figure 10 illustrates a proposed model for PEDV-induced immune responses, highlighting the overexpressed PRRs, APs, TFs, and IFN receptor-related genes that activate IFN and ISG signaling pathways in weaned piglets. Consistent with previous studies [23,31–33], type III IFNs, particularly IFN-λ3, and their receptor (IFNLR1), were generally expressed at higher levels than type I IFNs in all piglets. This result underscores the importance of type III IFNs in intestinal immunity and their antiviral role against PEDV compared to type I IFNs. Regardless, both IFNs lead to the production of various ISGs. Some of them, such as RIG-I/MDA5, TRIM25, IRF7, and STAT1, act as regulators of IFN signaling. The remaining ISGs function as antiviral effectors, obstructing viral replication by preventing viral entry and degrading viral ssRNA [19,21,44].

**Figure 10.**
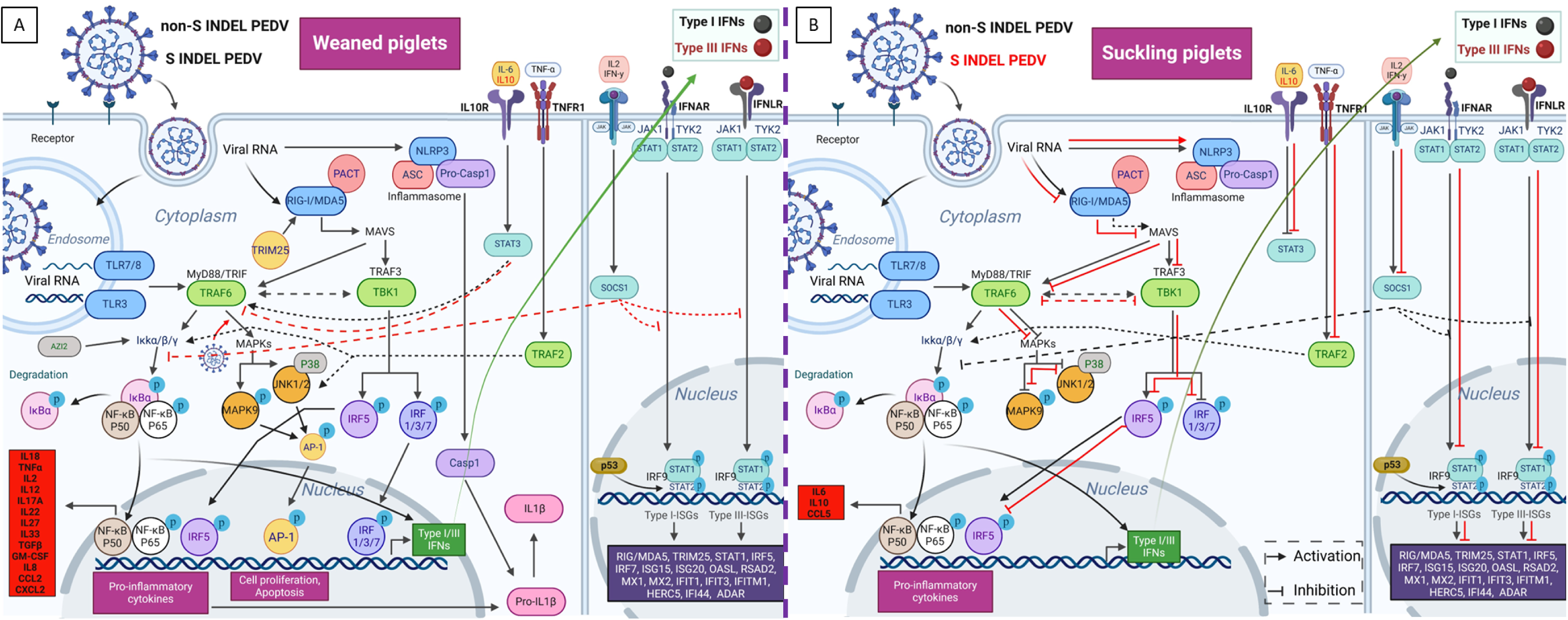
Proposed model for the diverse intestinal innate immune pathway responses induced by non-S INDEL and S INDEL PEDV strain in weaned (A) and ckling (B) piglets. The intestinal transcription pattern in weaned piglets is similar for both PEDV strains, whereas in suckling piglets, it differs between the n-S INDEL and S INDEL strains. In weaned piglets (A), the PRRs involved in PEDV recognition include TLR3, TLR7/8, RIG-I/MDA5, and NLRP3. The primary s involved in the recognition of PEDV infection through TLRs and RIG-I pathways are TRIM25/MAVS/IPS-1, TRAF6, and TBK1. These APs activate key TFse IRFs and NF-κB, which drive the transcription of antiviral genes and proinflammatory cytokines. IFN I and III synthesis in response to PEDV infection is ainly controlled by IRF7, IRF1, and NF-κB, with minimal involvement from IRF3, a crucial early regulator of type I IFNs. As a result, numerous ISGs are erexpressed through the JAK-STAT/IRF9 signaling pathway, along with other key ISGs and proinflammatory regulators like STAT3 and SOCS1. The primary ivers of proinflammatory responses to PEDV are NF-κB, IRF5, and IL-1β-mediated NLRP3 inflammasome activation, which also control the pyroptosis cell ath pathway via CASP-1 overexpression. Key regulators of cell survival, including MAPKs and the transcription factor AP-1, are also overexpressed llowing PEDV infection. Most of these pathways are activated in infected enterocytes of suckling piglets (B) exposed to the non-S INDEL strain (black rows), whereas they are suppressed in response to the S INDEL strain (red lines). Created with BioRender.com

In addition to IFNs and ISGs, heat shock proteins (HSPs) such as HSP 27, HSP 70 and HSP 90, also exhibit antiviral effects against PCoVs [45–49]. Based on our findings, the HSP 14 transcript was upregulated in weaned piglets exposed to any of the PEDV strains suggests an antiviral role for these proteins, indicating a potential avenue for developing novel antiviral therapies.

In terms of inflammatory response, weaned piglets exhibited a proinflammatory reaction to both PEDV strains, highlighted by a significant upregulation of Th1 and Th17 cytokines at 48 hpi. Conversely, newborn piglets had elevated levels of Th2 cytokines, including the anti-inflammatory IL-10. This result suggest that Th1 and Th17 responses (such as IL-22) may favor clinical protection during the acute phase in weaned piglets, while an anti-inflammatory response dominates in newborns [30]. Additionally, this study also shows a negative correlation between IL-10 and IFNs expression and clinical outcome. In this context, it is tempting to hypothesize that the lack of an effective inflammatory response in newborns, driven by IL-10, contributes to the more severe clinical outcome. Despite its known antiviral and mucosal protective role in promoting IL-22 overexpression, STAT3 may also be involved in a negative feedback loop between IL-10 and IFN signaling through an unknown mechanism [30,50].

Based on obtained results, NF-κB is likely crucial in initiating the antiviral proinflammatory response. However, the overexpression of inflammasome-related IL-1β and IL-18 as well as IRF5, a critical transcription factor for M1 macrophage polarization and Th1- Th17 responses [51,52], indicates a more complex response against PEDV in weaned piglets. IL- 18 activates and recruits cytotoxic T-lymphocytes (CTLs), NK cells, and macrophages, leading to increased levels of INF-γ and NF-κB [53,54]. The observed overexpression of IL-18 and INF-γ as well as IL-2 and TNF-α, in this study suggests that these cells play a crucial role in virus clearance in weaned piglets exposed to both PEDV strains, supporting the assertion that a cytotoxic inflammatory response plays a key role in countering PEDV during the acute phase of infection. In contrast, the absence of IL-18 in newborns following PEDV infection correlates with their limited presence of CTLs and NK cells in the intestinal mucosa [3]. Although further studies are needed, IL-18 synthesis may also be associated with STAT3/IL-22 activation [30].

Interestingly, NF-κB and IL-1β were expressed in weaned piglets at levels comparable to their main inhibitors, NFKBIA and IL-1RA (encoded by IL-1RN), respectively. These data suggest critical simultaneous regulation of inflammatory and anti-inflammatory pathways to prevent the progression of inflammatory disease [55,56].

Furthermore, the increased gene expression of prostaglandin E2 (PTGS2) observed in weaned piglets suggests that proinflammatory eicosanoid-derived mediators play an important role in defense against PEDV, similar to their role in immune responses to other coronaviruses, such as severe acute respiratory syndrome coronavirus 2 [57–61]. These results also propose potential new therapeutic targets for disease control. Additionally, chemokines like CCL2 seem to strongly counteract both PEDV strains as previously observed [25]. However, the expression of CCL5 differs from earlier findings [35], suggesting the need for further research to clarify its role in disease protection.

In line with a previous report [20,62,63], our findings suggest that PEDV may exploit kinase-dependent pathways, such as MAPK/AP-1 and JNK/p38, via TRAF2, TRAF6, and TBK1 to enhance viral replication during the acute phase of infection in weaned piglets, whereas such modulation was absent in newborn piglets. This could explain the similar viral loads observed in both weaned and newborn piglets. In addition to kinase-dependent signaling pathways, the current study provides evidence that PEDV can regulate cellular surveillance and key programmed cell death pathways such as apoptosis and pyroptosis, by modulating STAT3 as previously reported [64,65]. We also found that the overexpression of pyroptosis-related genes (NLRP3 and CASP-1) in weaned piglets indicates that the inflammasome regulates PEDV-infected enterocyte death via Gasdermin D (GSDMD)-mediated pyroptosis [49,66,67]. In contrast, the lack of this response in newborn piglets suggests that both PEDV strains suppress pyroptosis, potentially aiding viral replication as an immune evasion strategy [67]. Additionally, CASP-1 may trigger other cell death pathways, such as apoptosis [68]. Given that PEDV can influence apoptosis, ER stress, the unfolded protein response, and autophagy [18], further research is needed to investigate these pathways and their role in disease pathogenesis.

In conclusion, the present study enhances our understanding of age- and strain-dependent PEDV pathogenesis and early clinical outcomes, emphasizing the critical roles of IFNs and ISGs in innate antiviral immunity and Th1-Th17 mediated inflammatory response during acute phases, which protect weaned piglets in contrast to newborn animals that suffer from a more severe overt disease.

## ACKNOWLEDGMENTS

The authors thank the IRTA-Monells staff for the care and handling of the animals in the BSL2 facility, IRTA-CReSA staff and Montse Amenós from CRAG research center for their help and technical support. We would also thank Jianqiang Zhang and Darin M. Madson (Department of Veterinary Diagnostic and Production Animal Medicine, Iowa State University, Ames, IA, USA) for providing the non-S INDEL strain used in this study. We would also thank the UAB statistics service for their help in carrying out the statistical study.

## AUTHOR CONTRIBUTIONS

Conceptualization, J.V.-A. and J.S.; methodology, C.L.-F., E.C., N.N., M.P., R.L., K.S., H. V. and P.M.H.H.; formal analysis, C.L.-F., E.C., N.N., M.P., R.L., K.S., H. V. and P.M.H.H.; investigation, C.L.-F., K.S., P.M.H.H., J.V.-A. and J.S.; resources, J.V.-A. and J.S.; data curation, C.L.-F.; writing—original draft preparation, C.L.-F.; writing—review and editing, K.S., P.M.H.H., J.V.-A. and J.S.; supervision, J.V.-A. and J.S.; project administration, J.V.-A. and J.S.; funding acquisition, J.V.-A. and J.S. All authors have read and agreed to the published version of the manuscript.

## DATA AVAILABILITY STATEMENT

Original data files are available on request.

